# ISSRseq: an extensible method for reduced representation sequencing

**DOI:** 10.1101/2020.12.21.423774

**Authors:** Brandon T. Sinn, Sandra J. Simon, Mathilda V. Santee, Stephen P. DiFazio, Nicole M. Fama, Craig F. Barrett

**Affiliations:** Department of Biology and Earth Science, Otterbein University, 1 South Grove Street, Westerville, OH USA 43081; Department of Biology, West Virginia University, 53 Campus Drive, Morgantown, WV USA 26506; Institute for Sustainability, Energy, and Environment (ISEE), University of Illinois at Urbana-Champaign, 1201 W Gregory Drive, Urbana, IL USA 61801; Department of Biology, West Virginia University Institute of Technology, 410 Neville Street, Beckley, West Virginia, 25801; Genetic Immunotherapy Section, National Institute of Allergy and Infectious Diseases, National Institutes of Health, Bethesda, MD USA 20892

**Keywords:** ISSRseq, reduced representation sequencing, *Corallorhiza*, population genomics

## Abstract

1. The capability to generate densely sampled single nucleotide polymorphism (SNP) data is essential in diverse subdisciplines of biology, including crop breeding, pathology, forensics, forestry, ecology, evolution, and conservation. However, the wet-lab expertise and bioinformatics training required to conduct genome-scale variant discovery remain limiting factors for investigators with limited resources.
2. Here we present ISSRseq, a PCR-based method for reduced representation of genomic variation using simple sequence repeats as priming sites to sequence inter simple sequence repeat (ISSR) regions. Briefly, ISSR regions are amplified with single primers, pooled, used to construct sequencing libraries with a commercially-available kit, and sequenced on the Illumina platform. We also present a flexible bioinformatic pipeline that assembles ISSR loci, calls and hard filters variants, outputs data matrices in common formats, and conducts population analyses using R.
3. Using three angiosperm species as case studies, we demonstrate that ISSRseq is highly repeatable, necessitates only simple wet-lab skills and commonplace instrumentation, is flexible in terms of the number of single primers used, and can generate genomic-scale variant discovery on par with existing RRS methods which require more complex wet lab procedures.
4. ISSRseq represents a straightforward approach to SNP genotyping in any organism, and we predict that this method will be particularly useful for those studying population genomics and phylogeography of non-model organisms. Furthermore, the ease of ISSRseq relative to other RRS methods should prove useful to those lacking advanced expertise in wet lab methods or bioinformatics.

## INTRODUCTION

Reduced representation sequencing methods (RRS), in concert with high-throughput sequencing technologies, have revolutionized research in agriculture, conservation, ecology, evolutionary biology, forestry, population genetics, and systematics (Altshuler et al., 2000; Van Tassell et al., 2008; Davey et al., 2011; Narum et al., 2013; Lemmon and Lemmon, 2013; Meek and Larson, 2019). These methods generate broad surveys of genomic diversity by sampling only a fraction of the genome, therefore allowing for multiple samples to be sequenced simultaneously which reduces sequencing costs (Peterson et al., 2012; Franchini et al., 2017). While there exists a bewildering array of RRS methods (Campbell et al., 2018), the most commonly employed are various forms of restriction-site associated DNA sequencing (RAD; Baird et al., 2008) and genotyping-by-sequencing (GBS; Elshire et al., 2011). Both involve fragmenting the genome with one or more restriction enzymes, ligating adapters and unique barcode sequences, pooling, optionally size-selecting the fragments for optimal sequencing length, amplifying resulting fragments, and sequencing typically with Illumina short-read technology (Davey et al., 2011). Flexibility in these methods lies principally in the selection of one or more restriction enzymes; e.g. one popular method, double-digest RAD sequencing, uses both a ‘rare-cutting’ and ‘common-cutting’ enzyme (Peterson et al., 2012).

While methods such as GBS and RAD are increasingly commonplace, they may be technically challenging and economically infeasible for researchers who lack specific expertise in molecular biology, bioinformatics, and/or have limited access to expensive computational resources or sophisticated and often dedicated instrumentation. Thus, the need remains for simple and extensible methods for generating genome-scale variation. An alternative class of methods focuses on amplicon-based sequencing (Campbell et al., 2018, Ericksson et al., 2020). One such method is multiplexed ISSR genotyping by sequencing (MIG-seq; Suyama and Matsuki, 2015), which generates amplicons using primers comprising simple sequence repeat (SSR) motifs in multiplex, and sequences the inter simple sequence repeat (ISSR) fragment ends. Briefly, ISSR involves the use of primers that match various microsatellite regions in the genome (Zietkiewicz et al., 1994); the primers often include 1-3 bp specific or degenerate anchors to preferentially bind to the ends of these repeat motifs (Gupta et al., 1994; Zietkiewicz et al., 1994) or consist entirely of an SSR motif (Bornet and Branchard, 2001). MIG-seq uses ‘tailed,’ barcoded, ISSR-motif primers in a 2-step PCR protocol, first with the amplification of ISSR, and the second with common primers matching the tails to enrich these amplicons, followed by size selection and sequencing on the Illumina platform. This method has the advantage of not requiring more conventional Illumina library preparation (i.e. fragmentation of genomic DNA and ligation of barcoded sequencing adapters), as the adapters and barcodes are included in the tailed ISSR-motif primers, thus reducing the cost and labor associated with conducting many library preps.

While MIG-seq has been cited by more than 90 studies to date (e.g. Tamaki et al., 2017; Park et al., 2019; Gutiérrez-Ortega, 2018; Takata et al., 2019; Eguchi et al., 2020), many questions remain with regard to its efficiency and reproducibility. First, the use of long, tailed, ISSR primers raises the possibility of primer multimerization and unpredictability of binding specificity (eight forward and eight reverse primers in multiplex, as has typically been implemented). Second, as originally implemented, the protocol requires 96 unique forward-indexed primers, each 61 bp in length. The cost of synthesizing such lengthy primers equates to thousands of US dollars spent up-front, which may be prohibitive for many researchers, though these costs could be mitigated somewhat via dual indexing and the use of shorter adapter sequence tails. Third, MIG-seq only produces sequence data from the ISSR amplicon fragment ends, e.g. as is done with ribosomal DNA metabarcoding, potentially missing significant levels of variation by not sequencing the entire amplified fragments. The numbers of SNPs reported in studies using MIG-seq range from a few hundred in intraspecific studies (e.g. Suyama and Matsuki, 2015) to thousands in a species-level phylogenetic study (Eguchi et al., 2020), although missing data comprised the majority (81.4%) of the SNP matrix generated via the latter study. Data of this quantity and completeness can be useful for basic population genetics or phylogeographic studies, but are inadequate for studies requiring densely sampled polymorphisms across the genome (e.g. Quantitative Trait Locus mapping, genomic scans of adaptive variation, pedigree analysis). Indeed, Suyama and Matsuki (2015) state: “…the number of SNPs is fewer in our method [than in RAD-seq] (e.g. ∼1,000 vs. ∼100,000 SNPs), which means low efficiency in terms of the cost per SNP and sequencing effort.”

Here we present ISSRseq, a novel RRS method that is straightforward, extensible and uses single-primer ISSR amplicons. Briefly, our approach is to: produce single-ISSR primer amplicons (as opposed to using tailed, highly multiplexed primer pairs), pool amplicons from multiple primers per sample (Fig. 1), conduct low-cost, fragmentase-based Illumina library preparation, and sequence entire ISSR regions on the Illumina platform (Fig. 2). Further, we provide a user-friendly set of UNIX BASH scripts that together comprise an analysis pipeline for user-customized data quality control and analysis, output of SNP data in formats commonly used for population genomics and phylogeography, and an easy to use script template for downstream population genomic analyses in R. Our findings demonstrate that ISSRseq can generate comparable levels of variation to RAD or GBS, and orders of magnitude more variation than MIG-seq, while containing low levels of missing data. We present case studies of the utility of this method in *Populus deltoides* W. Bartram ex Marshall, a species with a relatively small and accessible reference genome, and two mycoheterotrophic orchid species with large, uncharacterized genomes: *Corallorhiza bentleyi* Freudenst. and *C. striata* Lindl. All lab research was carried out by undergraduate students, demonstrating that this method is amenable and accessible to those with relatively limited experience in molecular biology.

**Figure 1.** Critical steps of ISSRseq polymerase chain reaction (PCR), including the approximate time projected for library preparation of 48 samples with four primer sets. The optional gel verification step depicts three different ISSR primers for a single individual. PCR pool aliquot in Step 3 dependent on number of primer sets chosen to create library.

**Figure 2.** Critical steps of ISSRseq library preparation, including the approximate time projected for library preparation of 48 samples with four ISSR primers.

## MATERIALS AND METHODS

### Lab Methods

#### Sample Collection and DNA extraction

Our experimental design was chosen to evaluate the performance of ISSRseq at differing taxonomic scales. We selected *C. bentleyi* to determine if our method was informative below the level of species, *C. striata* to evaluate performance across putative species boundaries, and *P. deltoides* to test ISSRseq within a single accession of a species with a well-characterized genome. We collected 87 individuals from 28 *C. striata* localities, as in Fama et al. (2020, unpublished data) and Barrett et al. (2011; 2018; Table 1), and 37 individuals from six *C. bentleyi* localities. Samples of *C. striata* were collected from ten US states and two Canadian provinces (British Columbia and Manitoba, Table 1). Sampling of *C. striata* was focused on the western USA and in particular on California, where the taxonomic status of populations in this complex is uncertain (Barrett et al., 2011; 2018). Samples of *C. bentleyi* were collected in Allegheny, Bath, and Giles Counties, Virginia, USA; and from Monroe County, West Virginia, USA (Table 1). For both species of *Corallorhiza*, approximately 0.2 g of perianth or ovary tissue was removed with a sterile scalpel (so as not to include seed material) and DNA was extracted using a CTAB DNA extraction protocol, modified to 1/10 volume (Doyle and Doyle, 1987). A negative control reaction using ultrapure water and no template DNA was included for each set of single-primer reactions. We collected leaf tissue of *P. deltoides* accession WV94 from a plantation located at the West Virginia University Agronomy Farm (39.658889, -79.905278) in Morgantown, West Virginia (Macaya-Sanz et al., 2017). Genomic DNA was extracted using a DNeasy plant mini kit (Qiagen, Hilden, Germany; Cat. No. 69104).

**Table 1.** Sampling regions and localities for *C. striata* individuals and sampling counties and localities for *C. bentleyi* individuals. United States and Canadian provinces are abbreviated.

| Region/County | Sampling locality | Number of individuals |
| --- | --- | --- |
| *C. striata* |  |  |
| Coast Ranges Cascades | Ashland County, OR | 2 |
|  | Lane County, OR | 2 |
|  | Marin County, CA | 5 |
|  | Santa County, Cruz, CA (1) | 3 |
|  | Santa County, Cruz, CA (2) | 4 |
|  | Sonoma County, CA (1) | 2 |
|  | Sonoma County, CA (2) | 1 |
| Sierra Nevada | Fresno County, CA | 7 |
|  | Mariposa County, CA | 3 |
|  | Nevada County, CA | 1 |
|  | Placer County, CA | 5 |
| var. *striata* N USA | Cache County, UT | 5 |
|  | Idaho County, ID | 4 |
|  | Lewis and Clark County, MT | 5 |
|  | Lewis County, WA | 2 |
|  | Natrona County, WY | 4 |
|  | Skamania County, WA | 2 |
|  | Thompson-Nicola Regional District, BC | 2 |
|  | Winnipeg Region, MAN | 2 |
| var. *vreelandii* SW USA | Graham County, AZ | 4 |
|  | Otero County, NM | 4 |
|  | Ouray County, CO | 5 |
|  | Tooele County, UT | 5 |
|  | Utah County, UT | 4 |
| *C. bentleyi* |  |  |
| RG | Alleghany County, VA | 5 |
| CG | Bath County, VA | 14 |
| BS | Giles County, VA | 3 |
| OR | Giles County, VA | 5 |
| WR | Giles County, VA | 5 |
| PM | Monroe County, WV | 5 |

#### ISSR Primers and PCR conditions

Twenty-one ISSR primers were selected from the UBC Primer Set #9 (University of British Columbia, Canada; paper available on GitHub (www.github.com/btsinn/ISSRseq), with the addition of five unanchored primers designed by the authors (Table 2). We used only eight of these primers in our study of *C. striata* to test extensibility of our method with regard to the number of ISSR primers used to generate amplicons for sequencing. After several rounds of initial optimization, reactions were set up in 10 μl volumes with: 5 μl 2×Apex PCR Master Mix (Genesee Scientific, San Diego, California, USA; Cat. No. 42-134), 0.5 μl 5M Betaine (Fisher Scientific, Waltham, Massachusetts, USA; Cat. No. AAJ77507AB), 1.0 μl of each single primer at 10 μM starting concentration (one primer per reaction; Integrated DNA Technologies, Coralville, Iowa, USA), 2.5 μl nuclease-free ultrapure water, and 1.0 μl template DNA (diluted to 20 ng/μl prior to PCR with Tris-EDTA pH 8.0). PCR reactions were set up as single master mixes with individual ISSR primers and aliquoted to 96-well PCR plates. Conditions were as follows for each primer: 5 min at 95°C; 30 cycles of 95°C (30s), 50°C (30s), and 72°C (1m); and a final extension of 10m at 72°C. Caution was taken during PCR amplification to avoid human, microbial, or plant-based contamination; all lab surfaces were disinfected with 10% sodium hypochlorite solution prior to amplification. PCR products were visualized on 1% agarose gels to verify reaction success (see Fig. S1 for an example gel image).

**Table 2.** List of ISSR primers used in this study for the three test species. Numbers listed under ‘ISSR primer’ correspond to the UBC Primer Set 9 names (paper available on GitHub www.github.com/btsinn/ISSRseq).

| ISSR primer | Motif | *P. deltoides* | *C. bentleyi* | *C. striata* |
| --- | --- | --- | --- | --- |
| 813 | (CT)8T | X | X |  |
| 814 | (CT)8A | X | X |  |
| 815 | (CT)8G | X | X |  |
| 817 | (CA)8A | X | X |  |
| 820 | (GT)8T | X | X |  |
| 824 | (TC)8G | X | X |  |
| 826 | (AC)8C | X | X | X |
| 834 | (AG)8YT | X | X | X |
| 836 | (AG)8YA | X | X | X |
| 840 | (GA)8YT | X | X | X |
| 843 | (CT)8RA | X | X |  |
| 845 | (CT)8RG | X | X |  |
| 848 | (CA)8RG | X | X |  |
| 855 | (AC)8YT | X | X |  |
| 856 | (AC)8YA | X | X | X |
| 857 | (AC)8YG | X | X |  |
| 858 | (AC)8RT | X | X |  |
| 859 | (TG)8RC | X | X |  |
| 860 | (TG)8RA | X | X |  |
| 868 | (GAA)6 | X | X | X |
| 873 | (GACA)4 | X | X |  |
| caa5 | (CAA)5 | X | X | X |
| cag5 | (CAG)5 | X | X | X |
| gtt5 | (GTT)5 | X | X |  |
| gat5 | (GAT)5 | X | X |  |
| cat5 | (CAT)5 | X | X |  |

#### Library preparation and sequencing of ISSR amplicon pools

PCR products were pooled for individual accessions across all primer amplification reactions, and the remaining volume of original reactions were stored in a -80°C freezer as backups. Pooled PCR products were cleaned of excess PCR reagents via Axygen® AxyPrep FragmentSelect-I Kit (Corning, Tewksbury, MA, USA, Cat. No. MAG-FRAG-I-50) and two 80% ethanol washes on a magnetic plate. Cleaned PCR pools were quantified via Nanodrop spectrophotometry and diluted with TE buffer to 5 ng/μl. Library preps were conducted with the QuantaBio sparQ DNA Frag and Library Prep Kit (QuantaBio, Beverly, Massachusetts, USA, Cat. No. 95194-096), a relatively inexpensive, rapid library kit that uses a fragmentase to shear genomic or amplified DNA. The library preparation protocol is described in detail elsewhere (www.github.com/btsinn/ISSRseq). Briefly, we scaled library preparation volumes by half (thus a 96-reaction kit yields 192 preps). The sparQ Frag and Library Prep Kit allows fragmentation and end repair in a single step followed by Y-yoke adapter ligation (Glenn et al. 2019). We then performed a magnetic bead cleanup, total library amplification with barcoded Illumina iTru primers (Glenn et al. 2019), and a final bead cleanup; the bead to sample ratio was 1:1 for both cleanups. Fragmentase time was optimized in earlier trials, and consistently gave the best target library sizes for Illumina sequencing at 3.5 minutes (by contrast, for high molecular weight genomic DNA fragmentation is suggested to be set at 14 minutes). Three μl of each library was run on a 1% agarose gel with GeneRuler 1 kb Plus DNA Ladder (ThermoFisher Scientific; Cat. No. SM1331) followed by quantification via Qubit™ dsDNA BR Assay Kit (Invitrogen, Carlsbad, California, USA; Cat. No. Q32850). Amplicons for each accession were pooled at equimolar ratios. Size selection of an aliquot of the final pool was conducted using magnetic beads at a bead to sample ratio of 1:1, but users could also conduct two selective bead cleanups with different sample:bead ratios or gel excision. Fragment size range and intensity were quantified on an Agilent 2100 Bioanalyzer (Agilent Technologies, Santa Clara, CA, USA) and final library quantification was conducted via quantitative PCR at the West Virginia University Genomics Core Facility. The final library size ranged from 250-550 bp. Pooled, indexed ISSR amplicons were sequenced using 2 × 150 bp Illumina MiSeq (reagent kit v2) for *C. bentleyi* and *P. deltoides*. For *C. striata*, three sequencing runs were conducted: one lane each of 2 × 100 bp and 2 × 50 bp on a HiSeq1500 at the WVU-Marshall Shared Sequencing Facility, and one run of 2 × 150 bp (reagent kit v2) Illumina MiSeq at the WVU Genomics Core Facility.

#### Bioinformatics and Downstream Analyses

Our bioinformatic analysis pipeline consists of five BASH scripts for use on UNIX-based systems, each with customizable options, allowing simple but flexible parameter adjustment to fit the needs of a particular dataset or project (Fig. S2). We also supply an R script template to conduct various population genomic analyses. These scripts are not meant to dictate how ISSRseq data are treated or analyzed by users, but rather they are meant to serve as an accessible example of analysis potential for these data. Below, we detail the workflow iteratively and in the order of script usage. One of the BASH scripts, ISSRseq_ReferenceBased.sh, is optional if the user wishes to map to a reference genome, previously assembled contigs, or to a previously assembled set of ISSRseq amplicons. BASH and R scripts are provided and a detailed wiki with usage examples can be found via the ISSRseq GitHub repository wiki (www.github.com/btsinn/ISSRseq/wiki).

### 1) ISSRseq_AssembleReference.sh

#### Read Pre-processing

BBDUK (38.51, available from: https://sourceforge.net/projects/bbmap/files/) is used to remove adapters and priming sequences from reads, quality trim, and exclude GC-rich or poor reads for each sample. Additionally, we used the kmer trimming feature of BBDUK to remove ISSR motifs used as primer sequences from the ends of reads. Kmer length was set to 18 and the ‘mink’ flag set to 8. The use of the ‘mink’ flag allowed for the removal of matched kmers down to a length of 8 bp from that of the supplied priming sequences (Table 2). We also enabled trim by overlap (‘tbo’) and ‘tpe,’ which in tandem allowed for adapter trimming by leveraging read overlap and in such an event trimmed both read pairs to the same length to ensure adapter removal. Trimmed reads for which the average quality score was below 10 or length was less than 50 bp were excluded, with the exception of the HiSeq 50bp reads. Hard trimming of read ends was not conducted when combining MiSeq 100 & 150 bp, and HiSeq 50 bp data generated for *C. striata*.

#### Reference Assembly

We then used ABySS-pe (version 2.2.4, Jackman et al., 2017) to assemble the trimmed reads from the user-specified reference sample using a kmer length of 91. Kmer choice was guided by our desire to minimize potential assembly errors due to the presence of low-complexity repeats (SSR motifs). BBDUK was then used to trim the assembled contigs in the same fashion as was used for read trimming, but with the GC content filter set to 35% and 65% and with the entropy filter enabled and set at 0.85. A reference index and sequence dictionary were created by SAMtools ‘faidx’ (version 1.7-13-g8dee2a2, Li et al., 2009) and the Picard tool ‘CreateSequenceDictionary’ (version 2.22.8; Broad Institute), respectively.

#### Contaminant Filtering

Next, we used the trimmed and filtered reference contigs as queries in BLASTn (version 2.6.0, Camacho et al., 2009) to identify putative contaminant loci by using the human genome, and those of 153 genomes of organisms that could be expected as common contaminants of plant samples (see supplementary materials), as subjects with an e-value cutoff of 0.00001. Contigs with e-values below this cutoff are excluded from the final reference. The user also specifies a plastid genome, and/or any other genome of interest, to be used as a negative reference. Contigs that can be mapped to the negative reference by BBMap (version 38.51, available from: https://sourceforge.net/projects/bbmap/files/) were also excluded from the reference assembly. Contigs identified as putative contaminants or representing the negative reference are written to a FASTA file.

### 2) ISSRseq_CreateBams.sh

#### Read Mapping and BAM Creation

BBMap (version 38.51) was used to map trimmed reads from each sample to the assembled contigs using default mapping settings and killbadpairs enabled. SAMtools was used to sort and index the BAM (Cock et al., 2015) files of each sample. We then used the Picard tools MarkDuplicates (version 2.22.8; Broad Institute), to mark PCR and optical duplicate reads, and BuildBamIndex (version 2.22.8; Broad Institute) to re-index these final BAM files.

### 3) ISSRseq_AnalyzeBAMs.sh

#### Variant Calling and Filtering

We used the state-of-the-art variant calling pipeline GATK4 (version 4.1.8, McKenna et al., 2010) to call, filter, and jointly score variants amongst all samples simultaneously. HaplotypeCaller (Poplin et al., 2017) identified potential variant sites and called variants from locally-reassembled portions of each BAM, with --linked-de-bruijn-graph, --native-pair-hmm-use-double-precision, and -ERC GVCF modes enabled. GVCF files from each sample were then combined into a single VCF file with CombineGVCFs. GenotypeGVCFs was then used to perform joint variant scoring on the combined VCF file. Scored variants for downstream analysis were restricted to biallelic SNPs and INDELs, which were then hard-filtered guided by GATK Best Practices hard filtering recommendations (DePristo et al., 2011): “AF > 0.01 && AF < 0.99 && QD > 2.0 && MQ > 40.0 && FS < 60.0 && SOR < 3.0 && ReadPosRankSum > -8.0 && MQRankSum > -12.5 && QUAL > 30.0”. Since the workflow exists as a BASH script, users can customize any of the variant filtering parameters to suit their needs.

### 4) ISSRseq_CreateMatrices.sh

*Matrix Creation*. VCF2PHYLIP (version 2.0; Ortiz, 2019) was used to coerce the minor allele and hard-filtered SNP variants into nexus, phylip, and binary SNP formats with varying matrix inclusion thresholds corresponding to the minimum number of samples in which a variant was identified.

### 5 ISSRseq_ReferenceBased.sh [Optional]

This script processes input reads and prepares the necessary file structure for the use of the pipeline with a pre-existing reference, for example, if the user has a sequenced genome or previously generated *de novo* assembly of contaminant-filtered ISSR amplicons at their disposal. Unlike ISSRseq_AssembleReference.sh, this script does not conduct contaminant filtering or trim both ends of the sequence reads since the input reads are not used for *de novo* assembly of the reference. Users should remove organellar and other non-target contigs from the reference prior to using this script. The output directory can then be used for steps 2 - 4 outlined above.

#### Reference, coverage, and missing data comparisons for C. striata datasets

In order to test the effects of using different Illumina sequencing strategies (i.e. sequencing effort and read length), we collected three datasets for the same 87 accessions of *C. striata*: Illumina MiSeq 2 × 150 bp, HiSeq 2 × 50bp, and HiSeq 2 × 100 bp (see above). We were specifically interested in the total number of SNPs, mean coverage depth, and percent missing data for each of the three individual datasets plus a ‘combined’ dataset using all three. We mapped reads as above to a common reference based on the combined dataset, with the goal of producing the most complete reference for mapping reads from individual datasets.

#### Population Genetic Analyses in R

All population genetic analyses were carried out using packages in the R software environment (R Core Team 2019). We describe each of the analyses and the prerequisite data preparation and population strata used in the subsections below. An example R script is provided in the ISSRseq GitHub repository (www.github.com/btsinn/ISSRseq) and a walkthrough of these analyses is provided via the wiki (https://github.com/btsinn/ISSRseq/wiki/ISSRseq-R-Analyses).

##### Data preparation

Prior to population genetic analyses, we thinned our hard-filtered VCF files to retain one variant per locus by setting the *thin* flag of VCFtools (0.1.15; Danecek et al. 2011) to the maximum contig length, and then used the *max-missing* flag to remove variants for which missing data exceeded 90% and 80% for *C. bentleyi* and *C. striata*, respectively.

##### Analysis of Molecular Variance (AMOVA) and F-statistic Estimation

We used AMOVA to assess population structure by partitioning the genetic variation based on predefined population categories using the Poppr package (Kamvar et al. 2015; ‘*poppr*.*amova’* function). We performed a permutation test for 1,000 iterations to test phi statistics for significance (‘*randtest’* function). We also estimated commonly used population statistics including observed heterozygosity (H_o_), observed gene diversities (H_s_), inbreeding coefficient (F_is_), and fixation index (F_st_) across all loci using the hierfstat package (Goudet 2005; ‘*basic*.*stats*’ function) for both species. The hierfstat package was also used to estimate pairwise F_st_ (‘*genet*.*dist*’ function) among population categories.

##### Principal Components Analysis (PCA)

We used the adegenet package (Jombart & Ahmed 2011) to perform a principal components analysis (PCA) and discriminant analysis of principal components (DAPC) to assign sub-population membership probabilities to individual plants (‘*dapc*’ function).

## RESULTS

### Sequencing and Reference Assemblies

#### C. striata complex

PCR using eight SSR primers (Table 2) successfully generated amplicons from 81 of the 87 accessions. The total combined sequencing read pool comprised 310,379,211 read pairs, of which 239,142,938 remained after trimming and filtering (Table 3). We selected accession 253e_CA as the reference individual since it is the sample for which we recovered the greatest number of reads (28,935,452 read pairs). *De novo* assembly resulted in 17,143 contigs longer than 100 bp, comprising a reference assembly totaling 3,628,930 bp with an N50 of 212 bp and GC content of 47.24%. Putative contaminant filtering excluded 555 contigs that either mapped to a genome of a putative contaminant or to any plastome sequenced from the *C. striata* complex (JX087681.1, NC_040981.1, MG874039.1, NC_040978.1) or that of *C. bentleyi* (NC_040979.1). The reference assembly comprised 14,164 contigs shorter than 250 bp, 450 contigs longer than 500 bp, and 34 contigs longer than 1 kb; the longest contig was 2,032 bp. Contigs of putative contaminant or plastid loci totaled 144,068 bp with an N50 of 230 bp, GC content of 44.17%, and a maximum contig length of 1,399 bp. This reference assembly was used for all comparisons of variant scoring for *C. striata*, below. For sample 253e_CA, the average insert size was 290 bp, 65.6% of reads mapped to the final reference assembly, and 24.1% of reads mapped to the putative contaminant assembly. The mean coverage depth of filtered variants scored in accession 253e_CA using this reference was 123.29x.

**Table 3.** Comparison of the analysis of Illumina MiSeq 150 bp PE, and Illumina HiSeq 50 bp and 100 bp PE reads analyzed singly and in combination using the same de novo reference assembly for the *C. striata* complex. An asterisk denotes statistics which were calculated after the removal of six samples which produced few sequences. SNP = single nucleotide polymorphism; bp/SNP = Total base pairs of reference sequence per SNP; SD = standard deviation; # min = the minimum number of samples within which a SNP was scored to be included in the data matrix. Variant implies both INDELS and biallelic SNPs.

|  | **MiSeq 2 x 150** | **HiSeq 2 x 50** | **HiSeq 2 x 100** | **Combined** |
| --- | --- | --- | --- | --- |
| **# raw read pairs / # post-trim** | 9899895 / 13435422 | 151025536 / 97981594 | 149453780 / 138080666 | 310379211 / 239142938 |
| **Total HaploTypeCaller Variants** | 249697 | 328726 | 560669 | 641667 |
| **Filtered SNPs** | 8177 | 12694 | 28401 | 25904 |
| **Mean filtered variant QUAL score** | 1636.36 | 499.96 | 2982.01 | 4685.16 |
| **bp/SNP (total filtered SNPs)** | 410.42 | 280.03 | 122.10 | 140.10 |
| **Mean coverage depth/locus/accession** | 3.81 | 5.55 | 10.67 | 14.82 |
| **SD coverage depth** | 3.48 | 7.30 | 12.42 | 16.71 |
| **Mean coverage depth/locus/accession\*** | 4.09 | 5.94 | 11.44 | 15.88 |
| **SD coverage depth\*** | 3.44 | 7.41 | 12.52 | 16.82 |
| **SNPs/%missing data (min 1)\*** | 8177/54.8% | 12694/51.1% | 28401/42.7% | 25904/40.0% |
| **SNPs/%missing data (min 10)\*** | 7219/49.5% | 11808/47.9% | 26506/38.9% | 24168/35.9% |
| **SNPs/%missing data (min 50)\*** | 2412/20.7% | 3991/19.7% | 13387/18.2% | 13762/17.7% |
| **SNPs/%missing data (min 80)\*** | 261/3.1% | 627/2.3% | 3209/2.1% | 3503/2.1% |

#### C. bentleyi

Amplicons were successfully generated from 37 *C. bentleyi* accessions using 26 primers shown in Table 2. The total combined sequencing read pool comprised 40,949,418 read pairs, of which 25,523,858 survived trimming and filtering. Accession B8 was chosen as the reference individual, as it was the accession for which we recovered the greatest number of reads (913,462 read pairs). *De novo* assembly recovered 16,813 contigs longer than 100 bp, comprising a reference assembly totaling 3,928,602 bp in length with an N50 of 234 bp and GC content of 46.43%, excluding 686 contigs that either mapped to a genome of a putative contaminant or to the plastome of *C. bentleyi* (NC_040979.1). The majority of reference contigs were less than 250 bp in length (11,894), while 3,748 were longer than 250 bp, 1,003 were longer than 500 bp, and 168 were longer than 1 kb; the longest contig was 2,390 bp. Putative plastid or contaminant contigs totaled 259,774 bp with an N50 of 463 bp, GC content of 43.24%, and a maximum contig length of 2,436 bp. For sample B8, the average insert size was 421 bp, with 37.5% of reads mapped to the final reference assembly and 13.3% mapped to the putative contaminant assembly. The mean coverage depth of filtered variants scored in accession B8 using this *de novo* reference was 16.42x.

#### Populus deltoides WV94

We used all 26 SSR primers used in *C. bentleyi* to conduct PCR amplification of one *P. deltoides* clone (WV94), which was multiplexed along with *C. bentleyi* accessions. The raw sequencing pool comprised 589,403 read pairs, of which 390,992 survived trimming. Kmer trimming of adapter and/or SSR primer sequences occurred on 51.84% of reads. The chromosome-level assembly of *P. deltoides* (445, version 2.0; https://phytozome.jgi.doe.gov) comprises 403,296,128 bp. Trimmed reads covered 4,376,825 bp or 1.09% of the genome (55.96% of trimmed reads), covering a median of 1.04% (range = 0.62% to 1.85%; SD = 0.34) of each chromosome (Table S1). Visual observation of BAM files qualitatively suggested that SSRs were amplified relatively evenly throughout the genome (Fig. S3). A positive correlation between chromosome length and percent of each chromosome covered by mapped trimmed reads was recovered by linear regression (r^2^ = 0.412, p-value = 0.003; Fig. S4), congruent with our visual assessment of SSR amplification. Contrary to this positive correlation was read mapping to Chromosome 8, where 53,666 reads covered 1.85% of its total length, the largest of such values, despite its standing as the eighth longest chromosome. Mean coverage depth of filtered variants across all chromosomes was 36.71x.

### Read Mapping and Variant Calling Results

#### C. striata

Analysis of individual MiSeq and HiSeq sequencing runs of PCR amplicon pools and their combination resulted in variable numbers of raw variants, variant quality scores, variant coverage depth, and number and missingness of final filtered variants (Table 3). Exclusion of six samples, for which sequencing produced few reads, increased filtered variant depth for the MiSeq 150 run to 4.09 and that of the combined read pool to 15.88. Higher variant quality, coverage, and concomitant reduction in missingness of data guided our decision to use the analysis of the combined read pool for downstream analyses. 24,078 SNPs remained after VCFtools missingness filtering, and thinning to 1 variant per locus left 6,589 variants in the final *C. striata* matrix for analysis in R.

#### C. bentleyi

HaplotypeCaller identified 236,694 total variants. SNPs comprised 47,851 of filtered variants or 76.11 bp/SNP. The mean coverage and quality score of filtered variants across all samples were 9.75 and 314.13, respectively. Although the mean SNP quality score among the *C. bentleyi* samples was lower than that of SNPs scored from *C. striata*, missing data comprised only 10% of total filtered SNPs, 8.6% scored from at least 10 accessions, 7.0% scored from a minimum of 20 accessions, and 4.5% scored from 30 of the 37 accessions. After filtering with VCFtools, 51,132 variants remained in the *C. bentleyi* matrix and 3,536 variants remained after thinning for analysis in R.

#### P. deltoides WV94

Of the 8,134 variants called by HaplotypeCaller in the *P. deltoides* WV94 clone we sequenced, 1,040 SNPs passed hard filtering. Variants were identified on all chromosomes (Fig. S5), and all filtered variants were heterozygous.

#### AMOVA and population statistics

For the *C. bentleyi* dataset, which contained 3,536 variants, we found that 92.2% of genetic variation was explained within individuals which was significantly less than expected by chance (phi = 0.079, sigma = 23.1, p-value < 0.001), while 4.45% was explained among sampling localities within county which was significantly greater than expected by chance (phi = 0.046, sigma = 1.11, p-value < 0.002), and 3.44% of variation was explained among county (phi = 0.034, sigma = 0.860, p-value = 0.314) (Table 4). Average observed heterozygosity and observed gene diversity was 0.066 and 0.078, respectively (Table 5). The F-statistic coefficients for the *C. bentleyi* sampling locality grouped populations that had low inbreeding coefficients and genetic distances (Fig. 3a).

**Table 4.** Analysis of molecular variance (AMOVA) output with fixation index for *Corallorhiza* populations. # loci = number of variants analyzed; columns 3-5 are percent variation explained by each hierarchical level; ‘a/m regions-county’ = among regions (C. striata) or county (C. bentleyi); ‘a/m loc’ = among sampling localities within region or county; and ‘w/in ind’ = within individuals. Columns 6-8 are values of fixation index (Φ) and their significance: * P < 0.005, ** P < 0.001. “ΦCT” = among regions or county; “ΦSC” = among sampling localities within regions or county, and “ΦIT” = within individuals.

| Species | # variants | b/w<br>region-<br>county | b/w loc | w/in ind | $\Phi_{CT}$ | $\Phi_{SC}$ | $\Phi_{IT}$ |
| --- | --- | --- | --- | --- | --- | --- | --- |
| *C. bentleyi* | 3,536 | 3.44 | 4.45 | 92.2 | 0.034 | 0.046*** | 0.079** |
| *C. striata* | 6,589 | 49.2 | 8.24 | 42.5 | 0.492*** | 0.162*** | 0.575*** |

**Table 5.** *Corallorhiza* population statistics table for *C. bentleyi* sampling locality and *C. striata* region. # loci = number of variants analyzed, H_o_ = observed heterozygosity, H_s_ = observed gene diversities, F_is_ = inbreeding coefficient, F_st_ = fixation index.

| Species | # loci | $H_o$ | $H_s$ | $F_{is}$ | $F_{st}$ |
| --- | --- | --- | --- | --- | --- |
| *C. bentleyi* | 3,536 | 0.066 | 0.078 | 0.149 | 0.000 |
| *C. striata* | 6,589 | 0.077 | 0.096 | 0.199 | 0.198 |

**Figure 3.** Pairwise F_st_ values comparing **a**. *C. bentleyi* sampling localities and **b**. *C. striata* regions. Red indicates high relative differentiation.

For *C. striata*, we found that 42.5% of genetic variation was explained within individuals which was significantly less than expected by chance (phi = 0.575, sigma = 28.2, p-value < 0.001), while 8.24% was explained among sampling localities (phi = 0.162, sigma = 5.46, p-value < 0.001), and 49.2% of variation was explained among regions (phi = 0.492, sigma = 32.6, p-value < 0.001) which were both significantly greater than expected by chance (Table 4). Average observed heterozygosity and observed gene diversity was 0.077 and 0.096, respectively. The F-statistic coefficient values were moderately high for the *C. striata* when treating geographic region at the level of subpopulation (Table 5, Fig. 3b).

##### PCA results

We retained 3 PC axes for the *C. bentleyi* analysis which explained a total of 14.1% of the variation in the genetic dataset (Fig. 4a). Discriminant analysis of principal components revealed the number of clusters appropriate for assigning membership probability based on principal components was 2 for *C. bentleyi* (Fig. 4b). In the *C. striata* analysis we retained 4 PC axes which explained 32.0% of the total genetic variation (Fig. 4c). *C. striata* sample membership was best explained by four clusters (Fig. 4d).

**Figure 4.** Population structure analyses of *Corallorhiza bentleyi* (n = 37 accessions) and the *C. striata* complex (n = 81 accessions). **a**. adegenet Principal Components analysis (PCA) dotplot and **b**. DAPC membership probability barplot for *C. bentleyi* inferred from 51,132 variants. **c**. PCA dotplot and **d**. DAPC membership probability barplot for the *C. striata* complex inferred from 24,078 variants.

## DISCUSSION

### C. striata

Population genetic analyses of ISSRseq-generated SNPs correspond well to previously published phylogeographic patterns recovered using targeted sequence capture of plastid loci (Barrett et al., 2018) and population genetic estimates using nuclear markers (Barrett and Freudenstein, 2011). For example, on the basis of three nuclear introns Barrett and Freudenstein (2011) estimated mean Φ_CT_ as 0.450, using some of same accessions used in the present study, and we estimated the same parameter at 0.492 using 6,589 variants generated by ISSRseq. Furthermore, F-coefficients and the results of AMOVA and DAPC suggest the presence of population subdivision and geographic partitioning of genetic variation in like fashion with that found using previous AMOVA, STRUCTURE analyses (Barrett and Freudenstein, 2011) and phylogenomic inference (Barrett et al., 2018). The congruence of the results of independent analyses using thousands of SNPs identified using ISSRseq with those of previous studies suggests that these data found using our novel method: 1) are appropriate for commonly used analyses; 2) contain reliable population genomic signal; and 3) are preferable since ISSRseq generates genome-scale data without prior knowledge of the genome.

### C. bentleyi

ISSRseq corroborated results of a traditional ISSR investigation study of *C. bentleyi* conducted previously by our group. Fama et al. (2021) scored ISSR bands visually for two primers used in this study and found that 89% of molecular variance occurred within populations. Using ISSRseq, we likewise found that the majority of genetic variation was explained within sampling localities, with only 4.45% of the genetic variation explained by sampling locality. Additional evidence of the recovery of population genomic signal in these data was our identification of fixed homozygous sites exclusive to cleistogamous individuals. We find the general congruence of visually-scored ISSR banding patterns in Fama et al. (2021) with the more sensitive, sequence-based nature of ISSRseq to be additional corroboration of our new method.

### Populus deltoides WV94

Our analysis of a *Populus deltoides* WV94 clone demonstrated that the loci sequenced and variants identified by ISSRseq are located throughout the genome, and variants were not obviously clustered in particular chromosomes or their centromeres or telomeres (Figs S3 & S5). As expected, we observed that the depth of final filtered SNPs was greater (36.71x) than the median depth of coverage for each chromosome (1.04 x), which is to be expected with any reduced representation sequencing method.

#### ISSRseq is straightforward and extensible

ISSRseq generates genomic variants on a scale that is comparable to other established RRS methods, while minimizing time and wet-lab complexity. Assuming the availability of two PCR machines and familiarity with basic wet laboratory techniques, users can go from extracted DNA to an Illumina sequencing library for 48 samples and 4 primer sets within an eight-hour workday. This timeframe and capacity can be greatly increased for users who use 96 or 384 well plates to conduct PCR.

ISSRseq is suitable for users with minimal lab experience or resources, since only a thermocycler and commonplace equipment such as a DNA quantitation device and affordable neodymium magnets used for PCR cleanup with magnetic beads are necessary prior to sequencing. Indeed, the ISSRseq data analyzed here were generated by undergraduates, and ISSRseq projects have been conducted in an undergraduate course at West Virginia University. Additionally, the use of a commercially-available, fragmentase-based sequencing library preparation kit means that users can receive support directly from a company, rather than relying on personal communications with authors. While alternative library preparation protocols can be used with ISSRseq, we found the efficiency and reliability of a commercially-available kit that can prepare a library in about 2.5 hours to be fitting for our purposes. A sizable fraction of the cost of ISSRseq, as currently implemented, is the use of a library preparation kit. Our publication of this method stands as a proof of concept, rather than a prescription, and we expect that advanced users of ISSRseq will modify the protocol and analysis pipeline described herein.

The recovery of loci generated amongst samples sequenced using RAD-like RRS approaches is impacted by mutations to restriction sites, DNA sample impurities and degradation, and user error or random variation during size selection (Andrews et al., 2016). ISSRseq generates loci for downstream analysis via PCR rather than careful size selection of restriction-fragmented DNA via gel excision or Pulsed-Field electrophoresis, and we expect that locus dropout due to user error or DNA quality will prove to be minimal relative to RRS methods such as ddRAD. In line with this expectation, missing data accounted for only 10% of the 47,851 SNP data matrix we generated for *C. bentleyi* and only 4.5% missing data when we required a SNP to be called in 30 of 37 individuals. As a point of reference, Tripp et al. (2017) used RADseq to identify 302,987 SNPs in *Petalidium* spp. (Acanthaceae), however the number of SNPs included in their data matrices (1,568 to 53,792, depending on parameter choice) was only greater than that recovered using ISSRseq when the missingness cutoff was greater than 60%. While variable SNP recovery may be relatively less of a concern for phylogenetic applications (Eaton et al., 2017; Tripp et al., 2017; Piwczyński et al., 2020), they are known to negatively impact estimation of population genetic parameters. For example, missing data are known to bias estimation of effective population size (N_e_) and the inbreeding coefficient (F_is_; Marandel et al., 2020). The use of PCR for the generation of loci analyzed using ISSRseq may mean that locus dropout will be minimized across DNA samples of relatively lower concentration and/or molecular weight than is preferable when using many existing RRS methods.

#### Methodological Considerations, Suggestions, and Caveats

Not surprisingly, sequencing depth is an important consideration when using ISSRseq just as with using other RRS methods. Conducting fewer ISSR PCRs using diverse SSR motifs is one option to maximize the efficiency of sequencing depth and number of samples that can be multiplexed during sequencing. For example, we recovered similar cumulative assembled amplicon length in *C. striata* as *C. bentleyi* despite using eight PCRs and 26 PCRs for each, respectively. This is likely due to the fact that many of the primers used for *C. bentleyi* comprised similar SSR motifs but had different anchors, whereas we used fewer primers of differing SSR motifs in *C. striata*, which together reflect the diversity of SSR motifs amplified from the latter species. We also found that 100 bp, paired-end sequencing of the same library on a single HiSeq 2500 nearly tripled the mean sequencing depth per locus (4.09x vs 11.44x) and more than quadrupled the number of variants scored (8,177 vs 28,401) over a single lane of paired-end 150 bp sequencing on the MiSeq. Given these results, we suggest that users select primers representing diverse and dissimilar SSR motifs that produce the most amplicons and prioritize read number over read length. Furthermore, we recommend bead-based exclusion of the shortest PCR amplicons prior to library preparation to reduce sequencing of short amplicons likely comprising mostly low complexity sequence. Following these recommendations should also save on PCR reagent cost and worker time.

Although joint variant calling should be able to minimize false positives, we do recommend that future researchers consider conducting more stringent variant filtering that incorporates a control sequence to empirically determine variant filtering parameters. For example, GATK is able to leverage machine learning of known variants to recalibrate variant quality scores and determine appropriate filtering parameters for genomes that are already well characterized (DePristo et al., 2011). The implementation of more sophisticated variant filtration techniques will no doubt reduce noise in ISSRseq data.

The impact of phylogenetic distance or reference choice on locus recovery and variant calling was not tested in the current study. For example, phylogenetic distance between samples and a given reference genome has been shown to differentially impact variant scoring depending on the chosen genotype caller (Duchen and Salamin, 2020), a finding that certainly warrants additional study. In future applications of ISSRseq, users are encouraged to explore alternative SNP calling software or approaches for generating data matrices to be used in phylogenomic inference. Regardless, researchers are encouraged to use the BASH scripts provided as a template to guide them in designing a workflow that fits the needs of their particular study.

#### Potential Applications

In addition to studies of local adaption, population genomics, and phylogenetic inference, ISSRseq could prove a useful technique to those interested in RRS using longer loci, agricultural crop or livestock authentication, forensics, and microbiome studies. For example, the use of PCR to amplify loci for sequencing means that the theoretical maximum of locus length is limited only by the DNA polymerase used during the PCR step. By modifying the PCR conditions and reagents, users could conduct long range amplifications to recover loci that are thousands of base pairs in length. These long loci could facilitate robust estimates of linkage disequilibrium and allele phasing in species without pre-existing genomic resources, and for the construction of gene trees for phylogenomic studies, allowing the use of many multilocus coalescent analyses (ASTRAL, Bayesian species delimitation, etc.). Furthermore, due to the ubiquity of SSRs in genomes, ISSRseq could be used to conduct RRS for the investigation of microbiomes, symbioses, and even simultaneous sequencing of hosts and one or more pathogens.

## ACKNOWLEDGEMENTS

This work was funded by the WVU Department of Biology, the WVU Eberly College of Arts and Sciences, a WVU Program to Stimulate Competitive Research Grant to CFB, NSF Award 1920858 to CFB. NMF and MVS were supported by the West Virginia Research Challenge Fund through a grant from the Division of Science and Research, HEPC and in part by (i) the WVU Provost’s Office and (ii) the Eberly College of Arts and Sciences. SJS was supported by NSF award 1542509 to SPD. We thank the USDA Forest Service for permission to collect samples. We thank Ryan Percifield for technical assistance, and the WVU Genomics Core Facility for sequencing support and services, and CTSI Grant #U54 GM104942 which provides financial support to the WVU Core Facility.

## CONFLICT OF INTEREST

The authors are not aware of any conflicts of interest.

## AUTHORS’ CONTRIBUTIONS

CFB conceived ISSRseq. BTS, SJS, CFB, and SPD designed the laboratory and bioinformatic methodology and protocols. BTS, SJS, MVS, and NMF conducted laboratory work. CFB conducted all field work. BTS, SJS, CFB, and SPD analyzed the data. BTS, SJS, and CFB led the writing of the manuscript. All authors contributed to manuscript drafts and provided their consent for publication.

## DATA AVAILABILITY

Raw sequencing reads for all samples are available via NCBI BioProject PRJNA771539. Data matrices, scripts used and their usage instructions in wiki format, as well as the wet lab protocol, are available via GitHub (www.github.com/btsinn/ISSRseq) or Sinn et al. (2021).

## Supplementary Documents

Document of supplementary tables and figures

## SUPPLEMENTAL TABLES & FIGURES

**Supplementary Table 1.** Descriptive statistics of *P. deltoides* WV94 and mapping of trimmed reads.

| Chromosome | Length (bp) | Reads Mapped | bp covered | % bp covered |  |
| --- | --- | --- | --- | --- | --- |
| 1 | 53086243 | 52775 | 534218 | 1.006321 |  |
| 2 | 26748850 | 27372 | 166950 | 0.624139 |  |
| 3 | 22915696 | 16653 | 144066 | 0.6286783 |  |
| 4 | 24223889 | 24175 | 256558 | 1.0591115 |  |
| 5 | 26443402 | 28379 | 366857 | 1.3873291 |  |
| 6 | 28326727 | 41552 | 295106 | 1.0417935 |  |
| 7 | 16146797 | 5441 | 144415 | 0.8943879 |  |
| 8 | 21710561 | 53666 | 402354 | 1.8532639 |  |
| 9 | 13407431 | 30241 | 188690 | 1.4073539 |  |
| 10 | 21864811 | 16827 | 240152 | 1.0983493 |  |
| 11 | 18311809 | 9054 | 172726 | 0.9432492 |  |
| 12 | 15564500 | 19837 | 152948 | 0.9826721 |  |
| 13 | 17534440 | 22999 | 208538 | 1.1893052 |  |
| 14 | 17670024 | 11811 | 110966 | 0.6279901 |  |
| 15 | 15256731 | 20625 | 152503 | 0.9995785 |  |
| 16 | 14755466 | 12147 | 261082 | 1.7693918 |  |
| 17 | 16616056 | 3820 | 143324 | 0.8625633 |  |
| 18 | 17191025 | 31245 | 231940 | 1.3491924 |  |
| 19 | 15521670 | 9041 | 203432 | 1.3106322 |  |
| totals | 403296128 | 437660 | 4376825 | 1.0852633 | Avg. % covered |
|  |  |  |  | 1.0417935 | Median % covered |
|  |  |  |  | 0.3354717 | SD |

**Figure S1.**
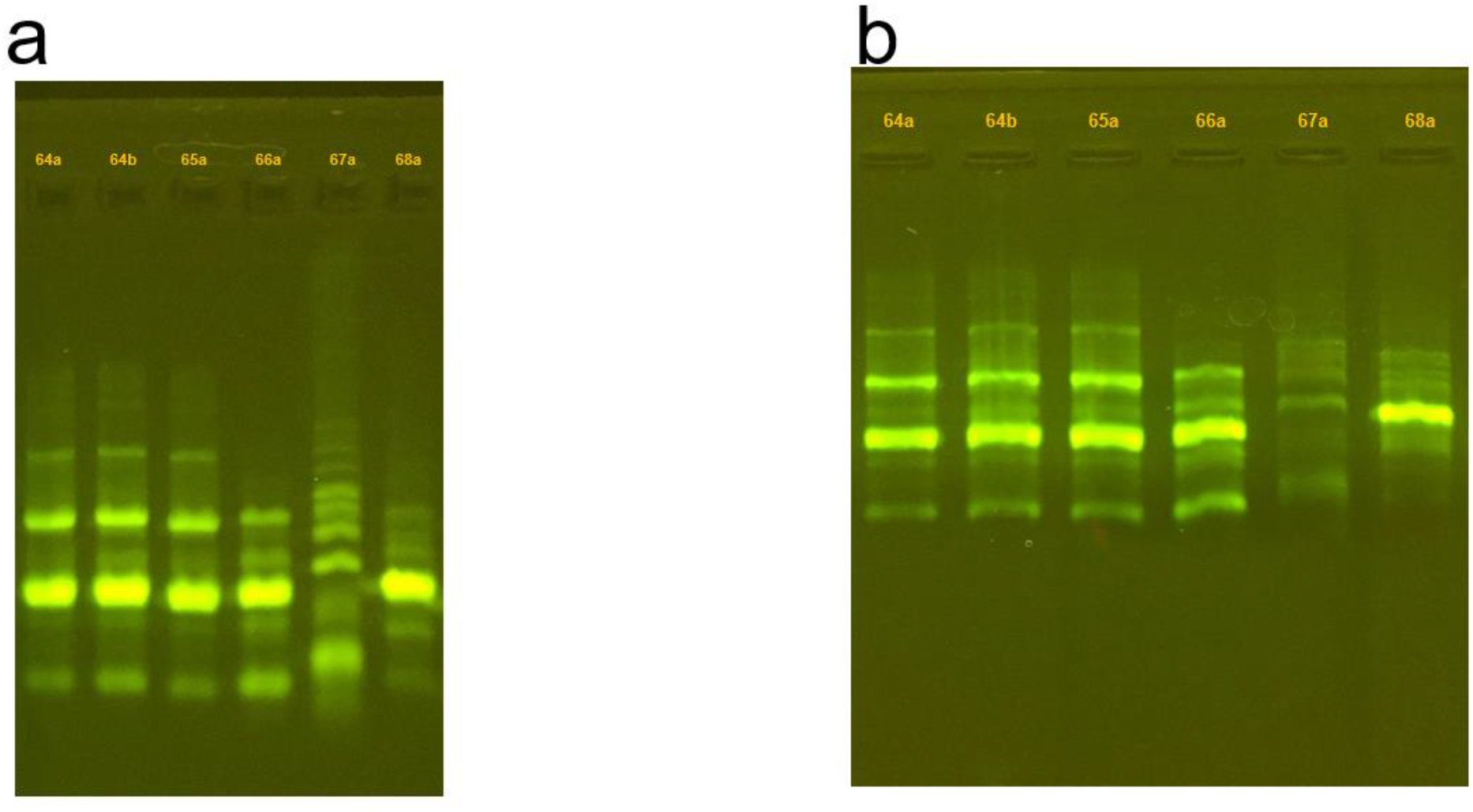
Gel electrophoresis (1% agarose) of PCRs conduced on A) 27 March and B) 31 May 2018 using ISSR Primer 868 on C. bentleyi samples: 64a, 64b, 65a, 66a, 67a, 68a (lanes labeled.) PCR products were allowed to separate differently by running gel A over a longer time period.

**Figure S2.**
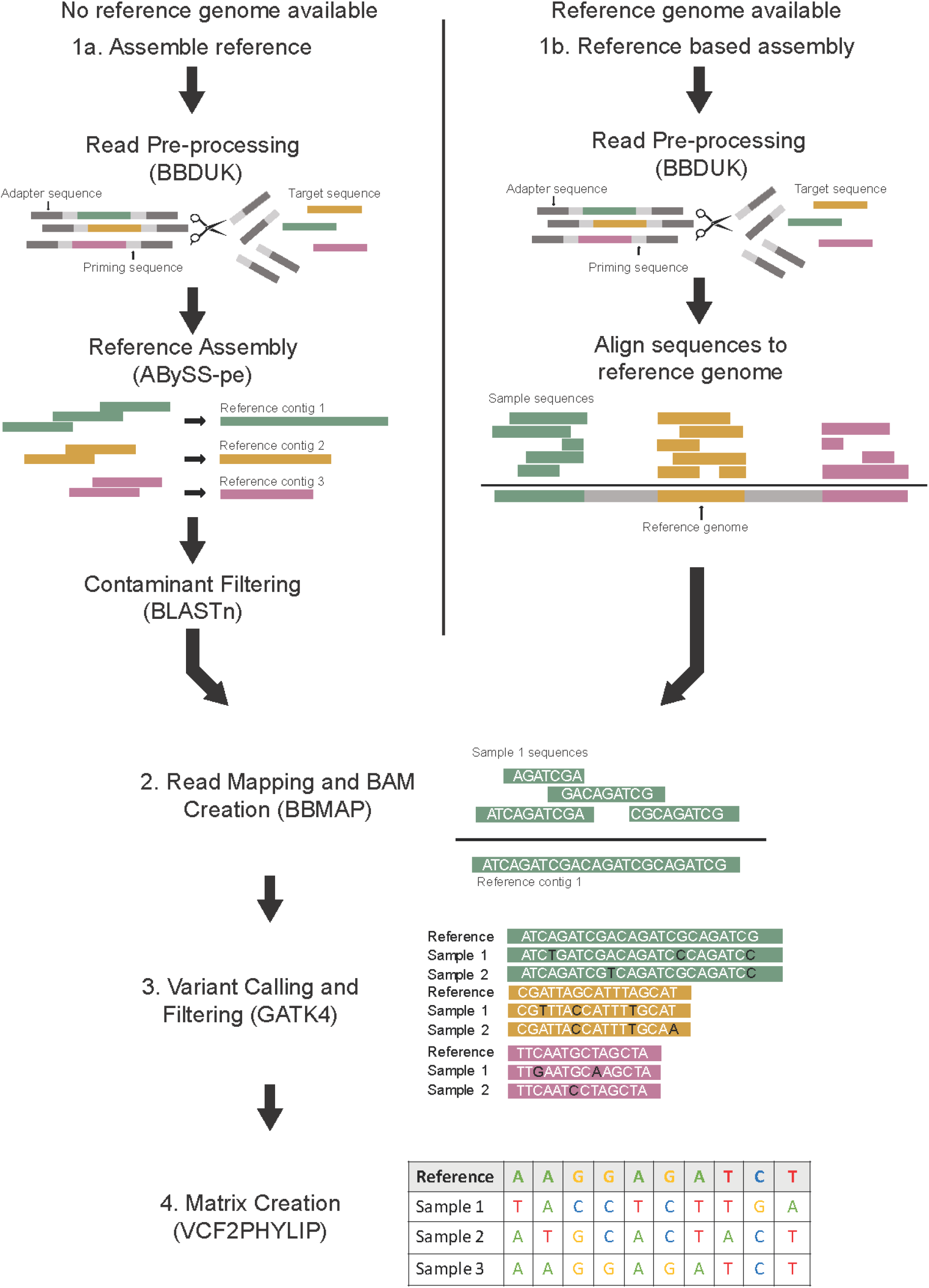
Flowchart for bioinformatic analysis pipeline with dependencies indicated in parentheses.

**Figure S3.**
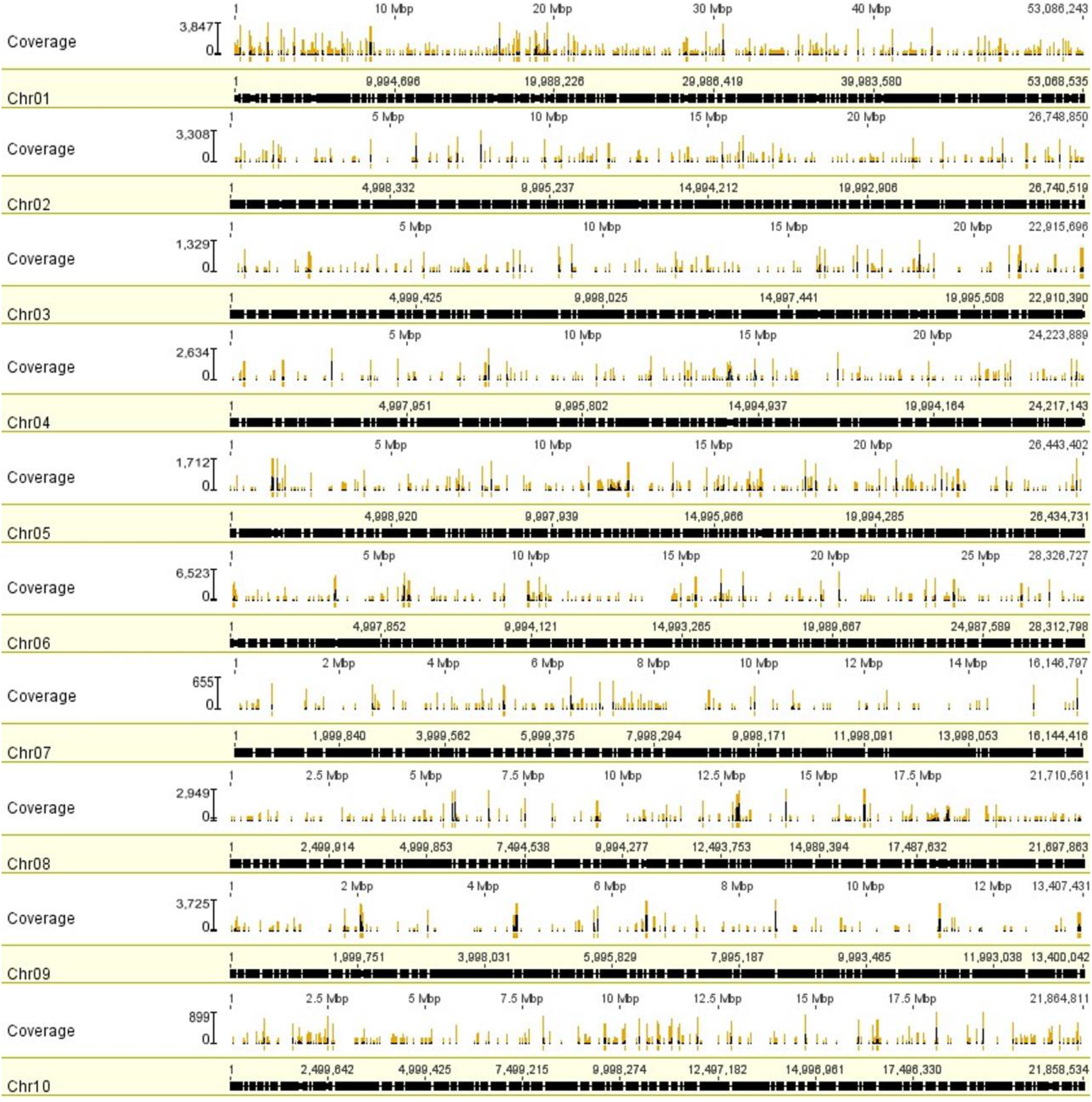
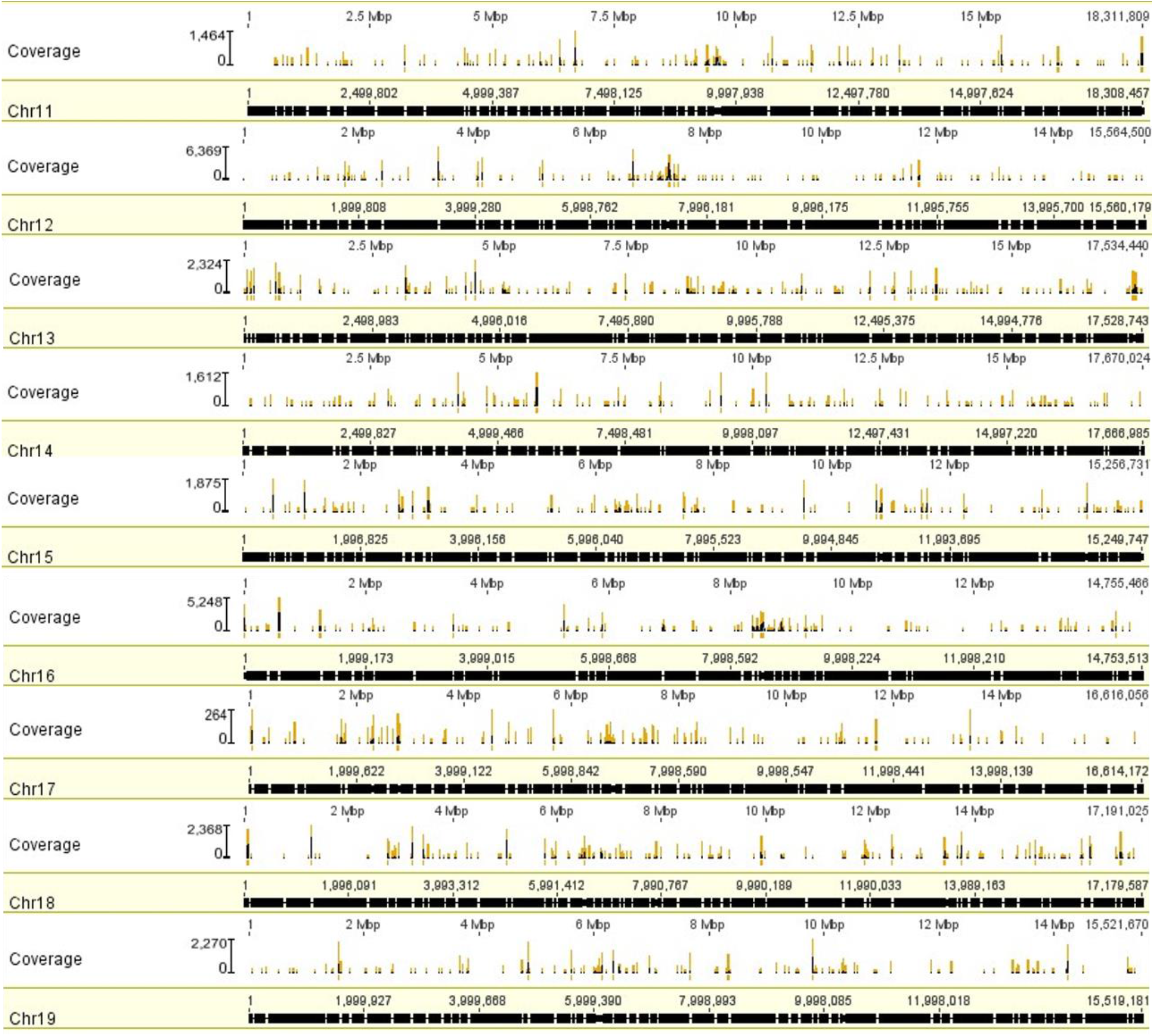
Mapping of trimmed reads to *P. deltoides* WV94 genome. Log-transformed coverage is shown above each chromosome, with regions of greater than 50x coverage depth denoted in yellow below the coverage histogram. Mapping of trimmed reads to *P. deltoides* WV94 genome. Log-transformed coverage is shown above each chromosome, with regions of greater than 50x coverage depth denoted in yellow below the coverage histogram.

**Figure S4.**
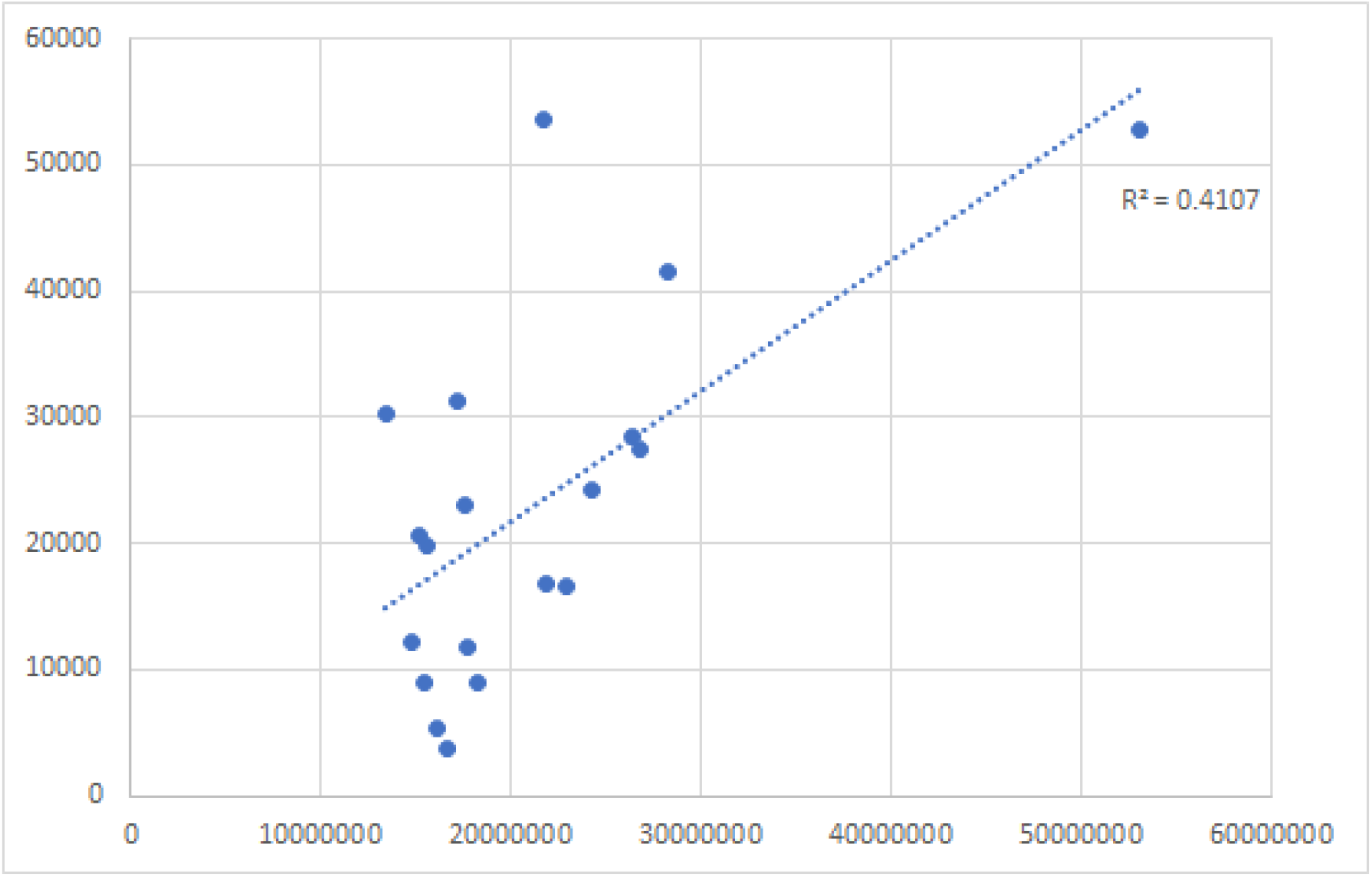
Trimmed reads mapped vs *P. deltoides* WV94 chromosome length. The dotted line represents the linear best fit (R^2^ = 0.4107; p-value = 0.003).

**Figure S5.**
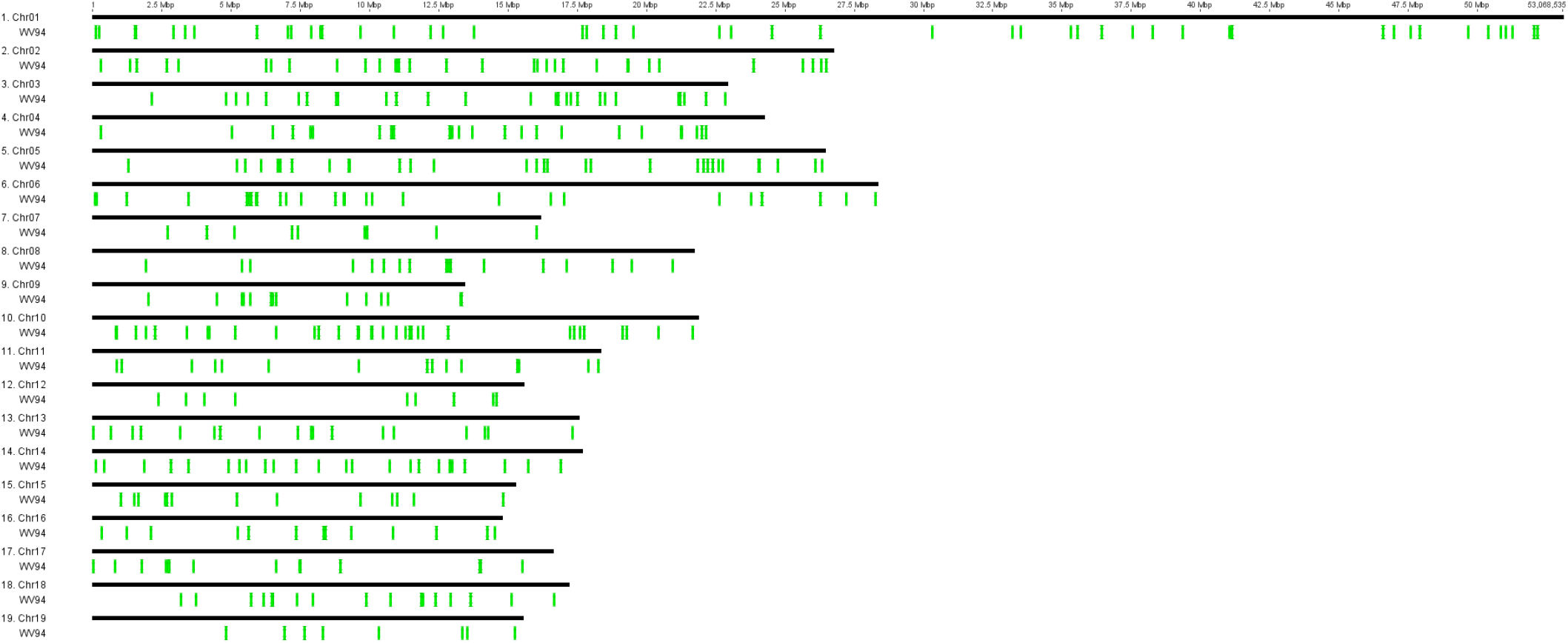
SNP variants annotated on the genome of *Populus deltoides* WV94. Variant positions are denoted with a green box below their position on each chromosome.

Supplementary File: Sources and identifiers of genomes used to filter putative contaminant loci.

Mycorrhizal Fungi (one from each genus available)

https://genome.jgi.doe.gov/portal/pages/dynamicOrganismDownload.jsf?organism=Mycorrhizal_fungi

*Acephala macrosclerotiorum* EW76-UTF0540 v1.0: Project: 1034881, Acema1

*Amanita muscaria* Koide v1.0: Project: 403202, Amamu1

*Boletus coccyginus* 2016PMI039 v1.0: Project: 1234182, Bolcoc1

*Cantharellus anzutake* C23 v1.0: Project: 1006245, Cananz1

*Cenococcum geophilum* 1.58 v2.0: Project: 402530, Cenge3

*Ceratobasidium* sp. anastomosis group I; DN8442 v1.0: Project: 1019533, CerAGI

*Choiromyces venosus* 120613-1 v1.0: Project: 1016739, Chove1

*Clavulina* sp. PMI_390 v1.0: Project: 1021554, ClaPMI390

*Cortinarius austrovenetus* TL2843-KIS7R v1.0: Project: 1193804, Coraus1

*Gautieria morchelliformis* GMNE.BST v1.0: Project: 1019425, Gaumor1

*Geopyxis carbonaria* DOB1671 v1.0: Project: 1127117, Geocar1

*Gigaspora rosea* v1.0: Gigro1

*Gyrodon lividus* BX v1.0: Project: 1016731, Gyrli1

*Hebeloma cylindrosporum* h7 v2.0: Project: 402607, Hebcy2

*Hyaloscypha finlandica* PMI_746 v1.0: Project: 1113379, Hyafin1

*Hydnum rufescens* UP504 v2.0: Project: 1019557, Hydru2

*Hysterangium stoloniferum* HS.BST v1.0: Project: 1019429, Hyssto1

*Kalaharituber pfeilii* F3 v1.0: Project: 1060175, Kalpfe1

*Kurtia argillacea* OMC1749 v1.0: Project: 1231371, Kurarg1

*Laccaria amethystina* LaAM-08-1 v2.0: Project: 1006253, Lacam2

*Lactarius akahatsu* QP v1.0: Project: 1177532, Lacaka1

*Lactifluus cf. subvellereus* BPL653 v1.0: Project: 1108981, Lacsub1

*Leucangium carthusianum* GMNB180 Co-culture: Project: 1191024, Leuca1

*Melanogaster broomeianus* MBLB.BST v1.0: Project: 1019457, Melbro1

*Meliniomyces bicolor* E v2.0: Project: 1008686, Melbi2

*Morchella importuna* CCBAS932 v1.0: Project: 1024000, Morco1

*Multifurca ochricompacta* BPL690 v1.0: Project: 1106235, Muloch1

*Oidiodendron maius* Zn v1.0: Project: 403613, Oidma1

*Paxillus adelphus* Ve08.2h10 v2.0: Project: 1006419, Paxru2

*Piloderma croceum* F 1598 v1.0: Project: 402979, Pilcr1

*Pisolithus albus* SI12 v1.0: Project: 1133208, Pisalb1

*Rhizophagus cerebriforme* DAOM 227022 v1.0: Rhice1

*Rhizopogon salebrosus* TDB-379 v1.0: Project: 1039526, Rhisa1

*Rhizoscyphus ericae* UAMH 7357 v1.0: Project: 1006265, Rhier1

*Russula brevipes* BPL707 v1.0: Project: 1106237, Rusbre1

*Scleroderma citrinum* Foug A v1.0: Project: 404724, Sclci1

*Sebacina vermifera* MAFF 305830 v1.0: Project: 403638, Sebve1

*Serendipita vermifera ssp. bescii* NFPB0129 v1.0: Project: 1053109, Sebvebe1

*Sphaerosporella brunnea* Sb_GMNB300 v2.0: Project: 1108957, Sphbr2

*Suillus americanus* EM31 v1.0: Project: 1052751, Suiame1

*Terfezia boudieri* ATCC MYA-4762 v1.1: Project: 1018931, Terbo2

*Thelephora ganbajun* P2 v1.0: Project: 1050999, Thega1

*Tirmania nivea* G3 v1.0: Project: 1108979, Tirniv1

*Tricholoma matsutake* 945 v3.0: Project: 404726, Trima3

*Tuber aestivum var. urcinatum* v1.0: Tubae1

*Tulasnella calospora* AL13/4D v1.0: Project: 404728, Tulca1

*Wilcoxina mikolae* CBS 423.85 v1.0: Project: 1016515, Wilmi1

*Xerocomus badius* 84.06 v1.0: Project: 1019549, Xerba1

Plant Pathogenic Fungi https://genome.jgi.doe.gov/portal/Plant_pathogens/Plant_pathogens.download.html

*Alternaria alternata* ATCC11680: Alalt1

*Ascochyta rabiei* ArDII: Ascra1

*Atropellis piniphila* CBS 197.64 v1.0: Project: 1134530, Atrpi1

*Blumeria graminis f. sp. hordei* 5874 v1.0: Project: 1145186, Bgh5874

*Botryosphaeria dothidea*: Botdo1

*Botrytis cinerea* v1.0: Botci1

*Cercospora zeae-maydis* v1.0: Project: 401984, Cerzm1

*Choanephora cucurbitarum* NRRL2744 (50) v1.0: Project: 1099013, Chocucu1

*Cladosporium fulvum* v1.0: Clafu1

*Cochliobolus carbonum* 26-R-13 v1.0: Project: 403760, Cocca1

*Colletotrichum abscissum* IMI 504890: Colab1

*Corynespora cassiicola* CCP v1.0: Project: 1019537, Corca1

*Cronartium quercuum f. sp. fusiforme* G11 v1.0: Project: 401997, Croqu1

*Cryphonectria parasitica* EP155 v2.0: Project: 16952, Cryphonectria

*Cytospora chrysosperma* CFL2056 v1.0: Project: 1110761, Cytch1

*Dactylonectria estremocensis* MPI-CAGE-AT-0021 v1.0: Project: 1103627, Daces1

*Diaporthe ampelina* UCDDA912: Diaam1

*Didymella zeae-maydis* 3018: Didma1

*Diplodia seriata* DS831: Dipse1

*Dothidotthia symphoricarpi* v1.0: Project: 1011345, Dotsy1

*Elsinoe ampelina* CECT 20119 v1.0: Project: 1064684, Elsamp1

*Elytroderma deformans* CBS183.68 v1.0: Project: 1166950, Elyde1

*Entoleuca mammata* CFL468 v1.0: Project: 1117716, Entma1

*Erysiphe pisi* Palampur-1 v2.0: Project: 1176372, Erypi2

*Eutypa lata* UCREL1: Eutla1

*Exobasidium vaccinii* MPITM v1.0: Project: 1016319, Exova1

*Fomitiporia mediterranea* v1.0: Project: 402518, Fomme1

*Fusarium fujikuroi* IMI 58289: Fusfu1

*Gaeumannomyces graminis var. tritici* R3-111a-1: Gaegr1

*Golovinomyces cichoracearum* UCSC1 v1.0: Project: 1055992, Golci1

*Gremmeniella abietina* DAOM 170408 v1.0: Project: 1104310, Greab1

*Grosmannia clavigera* kw1407: Grocl1

*Heterobasidion annosum* v2.0: Project: 16080

*Leptosphaeria maculans*: Lepmu1

*Macrophomina phaseolina* MPI-SDFR-AT-0080 v1.0: Project: 1103647, Macpha1

*Magnaporthe oryzae* 70-15 v3.0: Magor1

*Magnaporthiopsis poae* ATCC 64411: Magpo1

*Melampsora allii-populina* 12AY07 v1.0: Project: 1026279, Melap1finSC

*Microbotryum lychnidis-dioicae* p1A1 Lamole: Micld1

Microthyrium microscopicum CBS 115976 v1.0: Project: 1011369, Micmi1

*Mixia osmundae* IAM 14324 v1.0: Project: 404192, Mixos1

*Moniliophthora perniciosa* FA553: Monpe1

*Mycosphaerella graminicola* v2.0: Project: 16205

*Nectria haematococca* v2.0: Project: 16208, Necha2

*Neocosmospora boninensis* NRRL 22470 v1.0: Project: 1034400, Neobo1

*Neofusicoccum parvum* UCRNP2: Neopa1

*Neonectria ditissima* R09/05: Neodi1

*Oliveonia pauxilla* MPI-PUGE-AT-0066 v1.0: Project: 1103641, Olipa1

*Ophiostoma piceae* UAMH 11346: Ophpic1

*Paraconiothyrium sporulosum* AP3s5-JAC2a v1.0: Project: 1029422, Parsp1

*Phaeoacremonium aleophilum* UCRPA7: Phaal1

*Phaeomoniella chlamydospora* UCRPC4: Phach1

*Phakopsora pachyrhizi* MG2006 v1.0: Project: 1097278, Phapa1

*Phellinus igniarius* CCBS 575 v1.0: Project: 1108911, Pheign1_1

*Phoma multirostrata* 7a v1.0: Project: 1150440, Phomu1

*Phyllosticta citriasiana* CBS 120486 v1.0: Project: 1011301, Phycit1

*Pseudocercospora (Mycosphaerella) fijiensis v2*.0: Project: 16189

*Pseudoidium neolycopersici* MF1 Metagenome Minimal Draft: Project: 1145388, Psene1

*Puccinia coronata avenae* 12NC29: PuccoNC29_1

*Pyrenophora teres f. teres*: Pyrtt1

*Rhizoctonia solani* AG-1 IB: Rhiso1

*Saccharata proteae* CBS 121410 v1.0: Project: 1011317, Sacpr1

*Sclerotinia sclerotiorum* v1.0: Sclsc1

*Septobasidium* sp. PNB30-8B v1.0: Project: 1047727, Sepsp1

*Septoria musiva* SO2202 v1.0: Project: 401987, Sepmu1

*Setomelanomma holmii* CBS 110217 v1.0: Project: 1019737, Setho1

*Setosphaeria turcica* Et28A v2.0: Project: 401988, Settu3

*Sporisorium reilianum* SRZ2: Spore1

*Stemphylium lycopersici* CIDEFI-216: Stely1

*Taphrina populi-salicis* CBS419.54 v1.0: Project: 1166952, Tappo1

*Teratosphaeria nubilosa* CBS 116005 v1.0: Project: 1019765, Ternu1

*Testicularia cyperi* MCA 3645 v1.0: Project: 1040186, Tescy1

*Urocystis occulta* CBS 102.71 v1.0: Project: 1185590, Uroocc1

*Ustilago maydis* 521 v2.0: Ustma2_2

*Venturia inaequalis*: Venin1

*Verticillium alfalfae* VaMs.102: Veral1

*Zopfia rhizophila* v1.0: Project: 1011325, Zoprh1

*Zymoseptoria ardabiliae* STIR04_1.1.1: Zymar1

Human Genome

ftp://ftp.ensembl.org/pub/release-100/fasta/homo_sapiens/dna/

Homo_sapiens.GRCh38.dna_rm.primary_assembly.fa.gz

## Notes

https://github.com/btsinn/ISSRseq

### Competing Interest Statement

The authors have declared no competing interest.

